# Characterization of early white matter changes in CADASIL using microscopic diffusion imaging and relaxometry

**DOI:** 10.1101/2023.02.27.530286

**Authors:** David A. Barrière, Ivy Uszynski, Rikesh M. Rajani, Florian Gueniot, Valérie Domenga-Denier, Fawzi Boumezbeur, Cyril Poupon, Anne Joutel

## Abstract

**Background and purpose:** Cerebral small vessel diseases (SVDs) are characterized by early white matter (WM) changes, whose pathological underpinnings are yet poorly understood. CADASIL is a monogenic and archetypal SVD, providing an ideal model for investigating these changes. Here, we used multicompartment microscopic diffusion imaging and relaxometry to elucidate microstructural changes underlying early WM abnormalities in CADASIL.

**Methods:** We acquired diffusion MRI data with a multiple-shell Q-space sampling strategy, and relaxometry T1 and T2 data, with a 160 and 80-μm isotropic resolution respectively, *ex vivo*, in CADASIL and control mice. Diffusion datasets were computed with the Neurite Orientation Dispersion and Density Imaging model to extract the neurite density index, the extracellular free water and the orientation dispersion index. Relaxometry datasets were computed with a 3-compartment myelin water imaging model to extract the myelin content. MRI metrics were compared between CADASIL and control mice using voxel and WM tract-based analyses and with electron microscopy analysis.

**Results:** WM in CADASIL mice displayed a widespread reduction in general fractional anisotropy, a large increase in extracellular free water, a reduction in the myelin content, but no reduction in neurite density. Electron microscopy analysis showed a ∽2-fold increase in the extracellular spaces and an elevation of the g-ratio indicative of myelin sheath thinning in CADASIL WM.

**Conclusion:** Our findings suggest that accumulation of interstitial fluid and myelin damage are 2 major factors underlying early WM changes in CADASIL. Advanced diffusion MRI and relaxometry are promising approaches to decipher the underpinnings of WM alterations in SVDs.

## Introduction

Cerebral small vessel diseases (SVDs) account for up to a third of ischemic stroke and are a leading cause of cognitive impairment, dementia, disability and premature death occurring with aging. SVDs are driven by a complex mix of genetic and other risk factors, among which age and high blood pressure are deemed the most significant ^1^. CADASIL (Cerebral Autosomal Dominant Arteriopathy with Subcortical Infarcts and Leukoencephalopathy), the most frequent genetic SVD, is caused by dominant mutations in the gene coding for NOTCH3, a receptor predominantly expressed in small vessels ^2^. As a pure and archetypal SVD, CADASIL provides an ideal model to study disease mechanisms.

One consistent feature of SVDs is the presence of widespread white matter (WM) abnormalities that are detected as areas of WM hyperintensities by conventional brain magnetic resonance imaging (MRI) ^3^. It is currently hypothesized that WM alterations impair cognition by disrupting WM tracts ^4^. Diffusion tensor imaging (DTI), which measures diffusion properties of water molecules in tissues, can detect microstructural alterations of WM tracts. The typical pattern in patients with SVD is a reduction in fractional anisotropy (FA) and an increase in mean diffusivity ^5^. DTI has proved to be more sensitive than conventional MRI to detect WM changes and to better correlate with cognition ^6^. However, DTI provides only limited information about underlying changes in tissue properties, and pathological underpinnings of WM signal abnormalities are still poorly understood.

The development of new MRI techniques in combination with advanced biophysical models offers the opportunity to elucidate the microstructural changes underlying WM signal abnormalities. The use of multi-shell samplings of the diffusion Q-space allows it to go beyond the simple orientational profile of the diffusion process. It provides insight about the various diffusion processes occurring inside different cell populations and thus allows it to better consider the complexity of WM. When combined with multicompartmental models of brain tissue like the Neurite Orientation Dispersion and Density Imaging (NODDI) model ^7^, diffusion MRI (dMRI) allows us to map quantitatively geometric characteristics of cell populations within each voxel and provide metrics including the neurite (axons and dendrites) density, the neurite orientation dispersion index, and the content of extracellular free water ^8^. Using a similar approach, quantitative MRI (qMRI) relaxometry in combination with a 3-compartment modeling, like the myelin water imaging model, can provide direct and specific metrics of the myelin content ^9^.

In this study, we sought to explore microstructural changes underlying early WM abnormalities in SVDs. We used multicompartment microscopic diffusion imaging and relaxometry, *ex vivo*, in a well-established mouse model of CADASIL. We assessed the validity of our MRI findings using electron microscopy analysis.

## Methods

### Animals

Tg*Notch3*^R169C^ mice express the rat *Notch3* gene with the R169C mutation identified in CADASIL patients. As controls, we used Tg*Notch3*^WT^ mice, expressing the wild type rat *Notch3* gene and wildtype non transgenic (non-Tg) mice ^10^. In total, we used 13 Tg*Notch3*^R169C^, 12 Tg*Notch3*^WT^ and 11 non-Tg mice. All mice were maintained on an FVB/N background. Animals were housed under standard conditions (21–22°C; 12/12 h light/dark cycle) with food and water ad libitum. All experiments were approved by our local institutional Animal Care and Use Committees (CEEA9 and CEEA34, registration numbers 95 and 19-031) and by the French government (Ministère de l’Enseignement Supérieur et de la Recherche, authorizations #03653.02 and APAFiS #21100-2019041816189580v4).

### Preparation of samples for ex vivo MRI analysis

Mice were deeply anesthetized with sodium pentobarbital (80 mg/kg) and transcardially perfused with 0.1M heparinized phosphate buffer followed by 4% paraformaldehyde. After perfusion, animal heads were collected and post-fixed in 4% paraformaldehyde. Two days prior to the MRI session, the paraformaldehyde solution was replaced by a phosphate buffered saline solution. To avoid any possible damage and deformation, brains were kept inside the skulls during the entire procedure. For scanning, the head was placed into a custom-built MRI compatible tube filled with Fluorinert FC-40 (3M, Belgium), a magnetic susceptibility-matching artifact-free fluid which does not contribute to the NMR signal.

### Acquisition parameters

*Ex vivo* MRI data were acquired on a 11.7 Tesla BioSpec preclinical scanner (Bruker, Germany), with experimenter blinded to the genotype of animals. A Rapid Acquisition with Refocused Echoes (RARE) sequence was used to collect the series of T_1_-weighted volumes with the following parameters: RARE factor = 16, effective TE/TR = 8.37/2 500ms, field of view (FOV) = 19.2×15.2×15.2 mm^3^, acquisition matrix = 240×190×190, spatial resolution = 80×80×80 μm^3^, 1 average. This sequence (1h30min) was repeated with 32 different inversion times ranging from 150 to 2300 ms for a total scan duration of 44h16min. A multi-spin multi-echo pulse sequence was used to collect the series of T_2_-weighted volumes with the following parameters: TR = 510ms; 30 TE ranging from 5 to 155ms in steps of 5ms, FOV = 19.2×15.2×15.2 mm^3^, acquisition matrix = 240×190×190, spatial resolution = 80×80×80 μm^3^, 1 average, total acquisition time = 5h 10min. A multiple-shell diffusion-weighted MRI protocol was developed to sample the diffusion Q-space over 3 different shells and optimized for the NODDI model using a segmented 3D Pulsed Gradient Spin Echo EPI sequence with the following parameters: 9 segments, TE/TR = 24/250ms, FOV = 19.2×15.2×15.2 mm^3^, acquisition matrix = 120×95× 95, spatial resolution = 160×160×160 μm^3^, 1 average; diffusion sensitization b = 1,500/4,500/8,000 s/mm^2^ along 25/60/90 directions uniformly distributed over the unit sphere plus 3 reference volumes at b = 0 s/mm^2^, total scan duration 16h30min.

All data were pre- and post-processed using the Ginkgo toolbox freely available at https://framagit.org/cpoupon/gkg ^9,11–13^.

### Pre-processing of MRI data

MRI data were exploitable for 8 Tg*Notch3*^R169C^ mice (22.3 ± 2.2 months old), 7 Tg*Notch3*^WT^ mice (22.2 ± 2.3 mo) and 7 non-Tg mice (21.8 ± 1.8 mo). For 2 Tg*Notch3*^R169C^, 3 Tg*Notch3*^WT^ and 3 non-Tg mice, the metal ear tag had not been removed prior to the immersion of the head in fixative and MRI data were not exploitable. We used. Quantitative and diffusion-weighted MRI volumes were all co-registered to the Turone Mouse Brain Template and Atlas (60μm isotropic), which includes 1320 regions of interest ^14^ using the Advanced Normalization Tools ^15^. The corpus callosum was segmented using the Witelson and Aboitiz scheme segmentation ^15,16^.

### Post-processing of diffusion MRI data

The Generalized Fractional Anisotropy (GFA) map was calculated from the Q-space samples collected on the shell of largest b-value (b=8000s/mm^2^) using the analytical Q-ball model with a spherical harmonics order 6 and a Laplace-Beltrami regularization factor of 0.006 ^17^. To extract quantitative microstructural parameters, we used the NODDI microstructural model which relies on the partition of water diffusion in the brain into non-exchanging diffusive compartments ^7,18^. Within each voxel, the observed diffusion signal attenuation A results from the contribution of 4 different compartments with their own diffusion signal attenuation and volume fractions: the neurite compartment of water molecules trapped within axons and dendrites (A_neurite_, f_neurite_), the extra-neurite compartment of water molecules surrounding the neurites (A_extracellular_, f_extracellular_), the cerebrospinal fluid (CSF) compartment containing free molecules with an isotropic displacement probability (A_iso_,f_iso_) and an additional stationary compartment of water molecules trapped within spheres of nul radius assumed to correspond to glial cells (A_stat_,f_stat_).

### Post-processing of quantitative MRI data

The biophysical model used to assess the myelin water fraction (MWF) is based on the decomposition of the measured signals at the voxel level into components characterized by different relaxation times corresponding to different intra-voxel compartments ^9,19–21^. Within each voxel, we considered 3 different compartments corresponding to the myelin-related water, referred to as the myelin compartment, the unrestricted water referred to as the CSF compartment and the water related to both gray and WM. The 3-compartment model links the volume fraction of each compartment and their inherent relaxation times, with the T_1_-weighted measurements acquired for specific inversion times and the T_2_-weighted measurements acquired for specific echo times.

The signal to noise ratios from T_1_-weighted and T_2_-weighted relaxometry volumes were computed as the ratio between the mean signal from the brain parenchyma (brain mask) to the standard deviation of the noise evaluated on the image background. The estimation of noise levels with a standard deviation of respectively 0.003 and 0.04 for the T1-weighted and the T2-weighted volumes allowed us to approximate the non-centered chi-noise by a Gaussian noise, and thus to use non-negative least squares estimators ^9^ to compute quantitative T_1_and T_2_ relaxation times of the gray matter (T_1_=815.4±20.80ms, T_2_=48.22±2.11ms) and the WM (T_1_=775.9±19.34ms and T_2_=45.11±2.16ms) compartments. These qT1 and qT2 estimates were then used as initial parameters of our MWF mapper algorithm to calculate the volume fraction of each compartment and the proton density. A hybrid optimization scheme composed of a sequence of a Non-Linear Programming optimizer with constraints followed by a Monte-Carlo-Markov-Chain optimizer was implemented to robustly fit the MWF map.

All diffusion and quantitative maps were finally co-registered to the Turone Mouse Brain Template and Atlas using the previously calculated diffeomorphic transformations to perform statistics over control and mutant mice. Pre and post-processing of MRI data were performed with the experimenter blinded throughout the process.

### High-resolution electron microscopy analysis

We used 3 Tg*Notch3*^R169C^, 2 Tg*Notch3*^WT^ and 1 non-Tg mice aged 22 months. Mice were deeply anaesthetized and transcardially perfused with 0.1M heparinized phosphate buffer followed by Karlsson-Schultz fixative (4% paraformaldehyde, 2.5% glutaraldehyde in 0.1M phosphate buffer). Samples containing the most anterior part of the corpus callosum (genu) were processed using high pressure freezing and freeze substitution as previously described ^22^. For assessment of pathology, randomly selected, non-overlapping images of ultrathin sections were taken at 6,500× magnification. Quantification was performed as recently described, with the experimenter blinded throughout the process ^22^. G-ratios were corrected for inner tongue diameter and calculated as: [axon diameter]/([axon+myelin diameter]-([axon+inner tongue diameter]-[axon diameter])).

### Statistical analysis

To assess regional differences at the voxel level of dMRI, NODDI and MWF metrics, between controls and TgNotch3^*R169C*^ mice, smoothed maps were compared using a two-sample t-test implemented in Statistical Parametric Mapping 8 for voxel-wise comparison. The significance level applied was set at *p* = 0.05 (t(20) = 1.7274 false discovery rate (FDR)-corrected), with a minimum cluster size set at 25 voxels. From maps generated by the analyses and the mouse brain template, we made brain plots with the <u>xjView plugin</u> (https://www.alivelearn.net/xjview/). To assess the amplitude of differences within each of the major WM tracts, we performed a region of interest analysis. Clusters revealed by the voxel-wise comparisons were identified using the mouse brain atlas and an in-house pipeline developed with MATLAB Simulink 10b (The Mathworks, USA). Clusters identified were binarized and regions of interest obtained were used to extract each metrics with the <u>REX plugin</u> (https://web.conn-toolbox.org/). Statistical significance was determined using a 2-way ANOVA followed by a two-stage linear step-up method of Benjamini, Krieger and Yekutieli to correct for multiple comparisons by controlling the False Discovery Rate (<5%) (GraphPad Prism 9 software). Significance for electron microscopy data was calculated using an unpaired Student’s t test (GraphPad Prism 9 software).

## Results

### Design of the MRI study

We used the Tg*Notch3*^R169C^ line, a well-characterized mouse model of CADASIL, in which mutant mice aged 20 to 22 months exhibit asymptomatic WM abnormalities in the absence of lacunar infarcts ^10^. Therefore, we assumed that at this age mutant mice recapitulate the early stage of WM changes in patients. We collected in the same animals one dMRI dataset with a multiple-shell Q-space sampling strategy and two T1-weighted and T2-weighted qMRI relaxometric datasets at 11.7T. The dMRI and the relaxometry data sets were acquired with 160-μm and 80-μm isotropic resolution respectively over 3 consecutive days (Fig 1A). Figure 2 shows representative whole brain tractograms highlighting our ability to accurately delineate WM fiber tracts at ultra-high resolution in the mouse brain.

**Figure 1:**
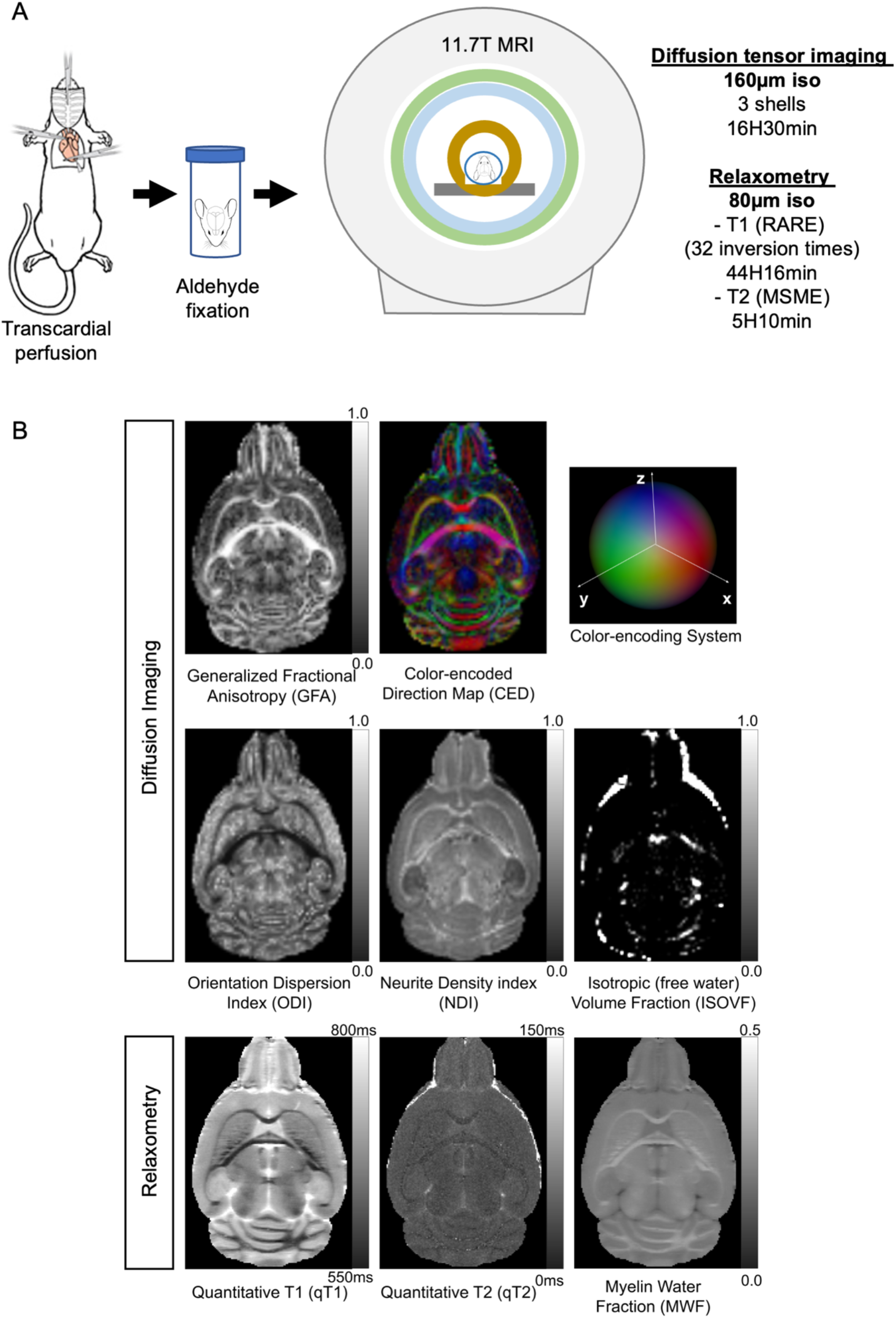
MRI paradigm to explore WM microstructural lesions in CADASIL and control mice. (A) Overview of the pipeline for *ex vivo* MRI analysis of mouse brain. The aldehyde fixed head of each animal was scanned over 66 hours. (**B**) Representative axial maps of dMRI metrics (GFA and directional brain map) (top row), NODDI metrics (ODI, NDI and ISOVF) (middle row) and quantitative relaxometry metrics (T1, T2 and MWF) (bottom row) in a control mouse.

**Figure 2:**
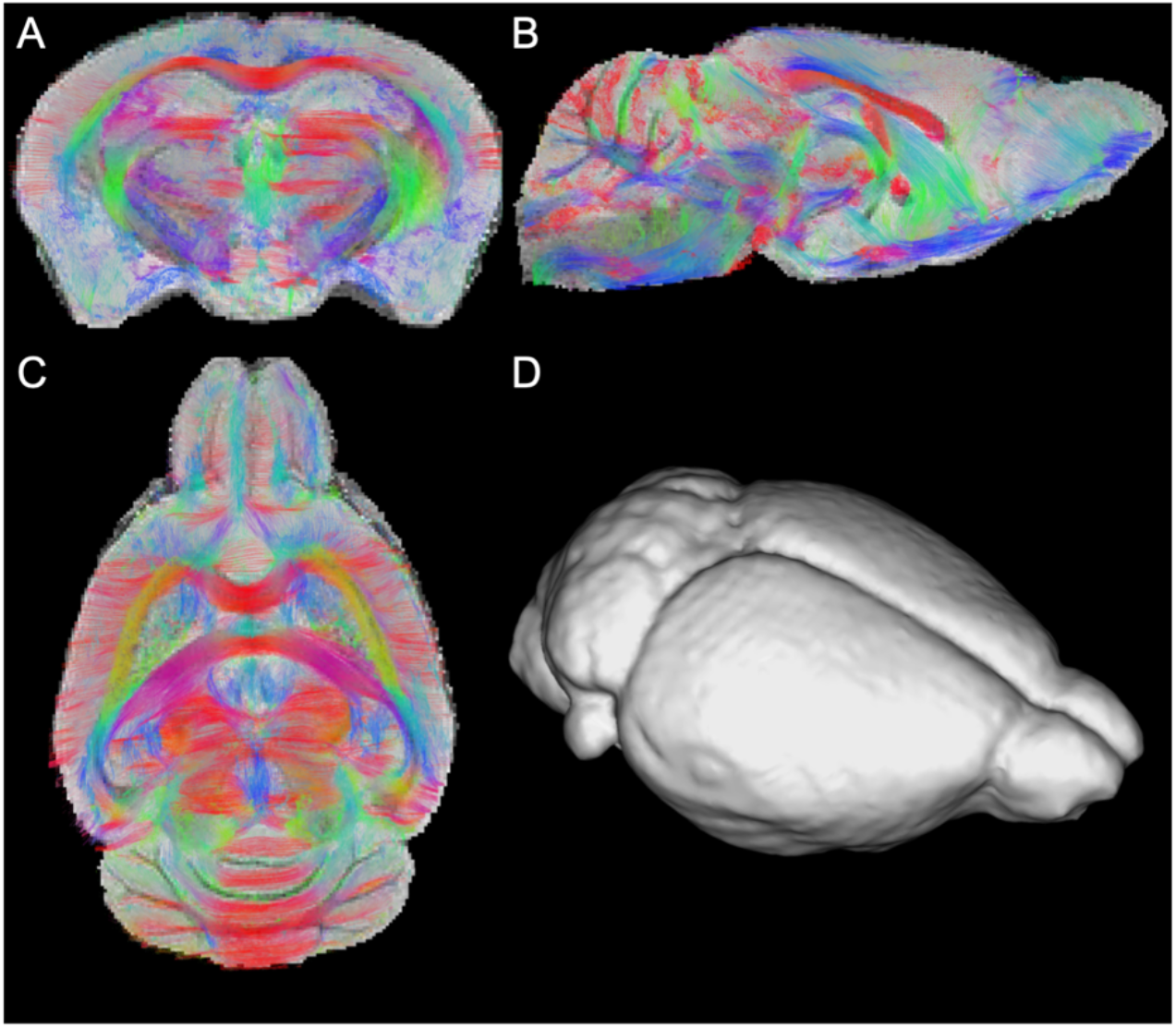
Color coded tractography reconstructions of WM tracts. Shown are coronal (**A**), sagittal (B) and axial (**C**) views and (**D**) a brain mesh.

From dMRI, we calculated the Generalized Fractional Anisotropy (GFA), a sensitive proxy of the loss of WM tract integrity. To probe the nature of microstructural abnormalities, dMRI data sets were then processed with the NODDI model to extract the neurite (axons and dendrites) density index (NDI), the isotropic volume fraction (ISOVF) corresponding to the extracellular free water and the orientation dispersion index (ODI). The relaxometry datasets were computed using a 3-compartment biophysical model to quantify the myelin water fraction (MWF), a direct metric of the myelin content. From these analyses, we obtained quantitative maps that were co-registered to a common template (Fig 1B).

### GFA

We focused our analysis on the 6 major hemispheric WM tracts in the mouse: the anterior commissura, the corpus callosum, the internal capsula, the optic tract, the fimbria hippocampi and the fornix. Non-Tg and Tg*Notch3*^WT^ mice showed comparable GFA in each WM tract and were pooled thereafter into a single control group for all analyses (not shown). GFA was widely reduced in WM tracts of Tg*Notch3*^R169C^ mice compared to control mice (Fig 3A-B). The corpus callosum, which is the largest WM tract in the mouse, was further segmented into 5 distinct regions including the anterior forceps, the genu, the body, the splenium and the posterior forceps. Generalized FA was reduced in all 5 regions in mutant mice (Fig 3C-D).

**Figure 3:**
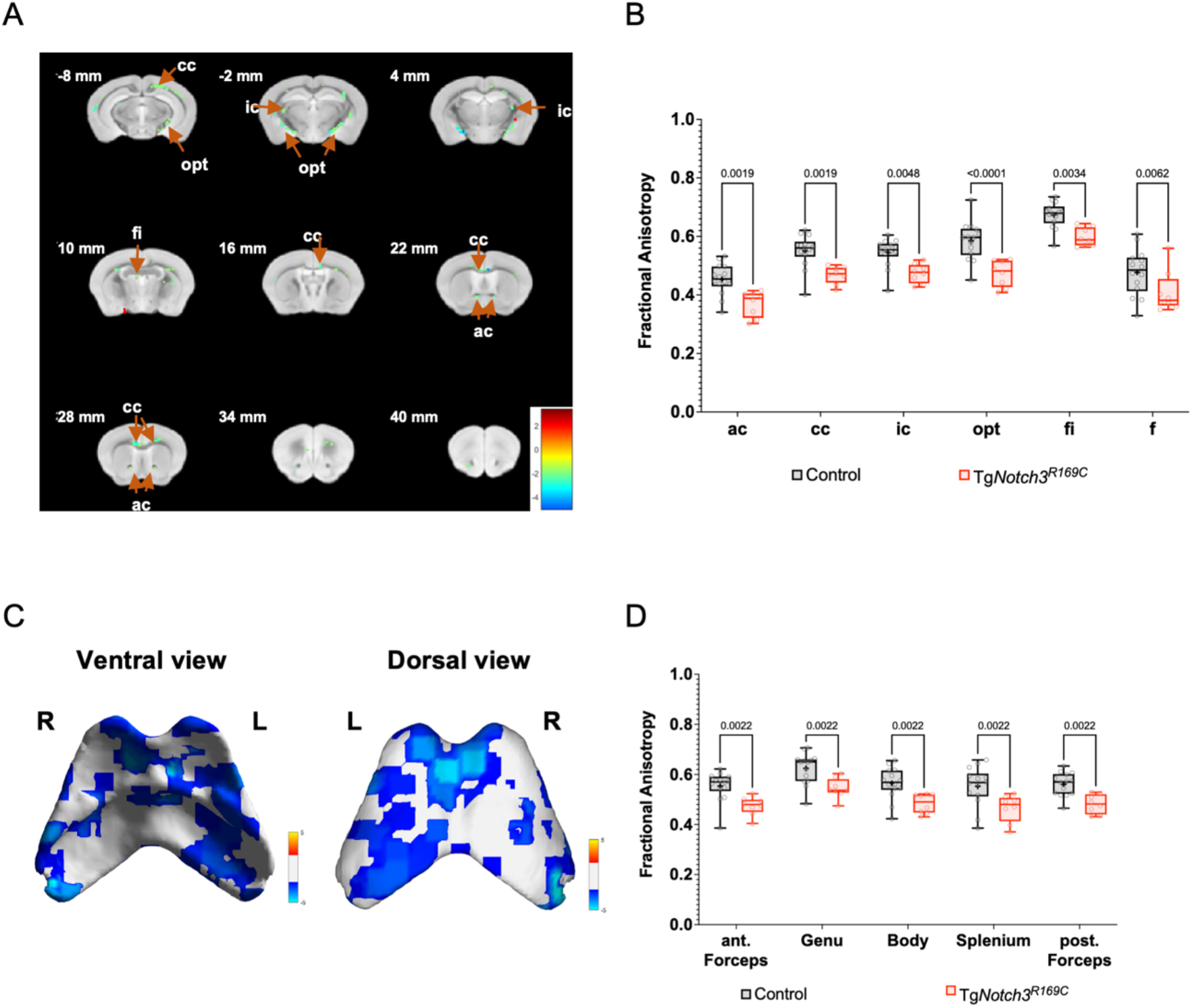
GFA is widely reduced in hemispheric WM tracts of CADASIL mice. (**A)**. Differences in voxel-based GFA values in WM tracts between Tg*Notch3*^R169C^ and control mice. Shown are clusters (threshold = 25 pixels) with values significantly different in Tg*Notch3*^R169C^ mice compared to control mice, overlaid on a series of coronal slices ranging from -6mm to +40 mm, with an interval of 6 mm. The color bar indicates the range of differences between mutant and control mice (**B**) Box-and-whiskers plots of quantitative values in the specified WM tracts of Tg*Notch3*^R169C^ and control mice analyzed by a two-way ANOVA followed by a two-stage linear step-up procedure of Benjamini, Krieger and Yekutieli to correct for multiple comparisons. Shown are the q values. ac, anterior commissura; cc, corpus callosum; ic, internal capsula; opt, optic tract; fi, fimbria of the hippocampus and f, fornix. (**C**) Differences in voxel-based GFA values in the corpus callosum between Tg*Notch3*^R169C^ and control mice. Shown are clusters (threshold = 25 pixels) with values significantly different in Tg*Notch3*^R169C^ mice compared to control mice, overlaid on ventral and dorsal views of the corpus callosum. (**D**) Box-and-whiskers plots of quantitative values in the specified corpus callosum regions of Tg*Notch3*^R169C^ and control mice.

### NODDI and MWF metrics

The NDI was essentially unchanged in WM tracts of Tg*Notch3*^R169C^ mice compared to controls, except in the internal capsule and the fornix where it was slightly increased (Fig 4A-B). The ISOVF (extracellular free water) was highly increased in all 6 WM tracts of mutant mice (Fig 4C-D). The ODI was also markedly increased in these WM tracts (Fig 4E-F). The MWF was reduced in all 6 WM tracts in Tg*Notch3*^R169C^ mice in comparison with control mice (Fig 5). In the corpus callosum, the ISOVF and ODI were widely increased and the MWF was reduced in mutant mice compared with control mice (Fig 6).

**Figure 4:**
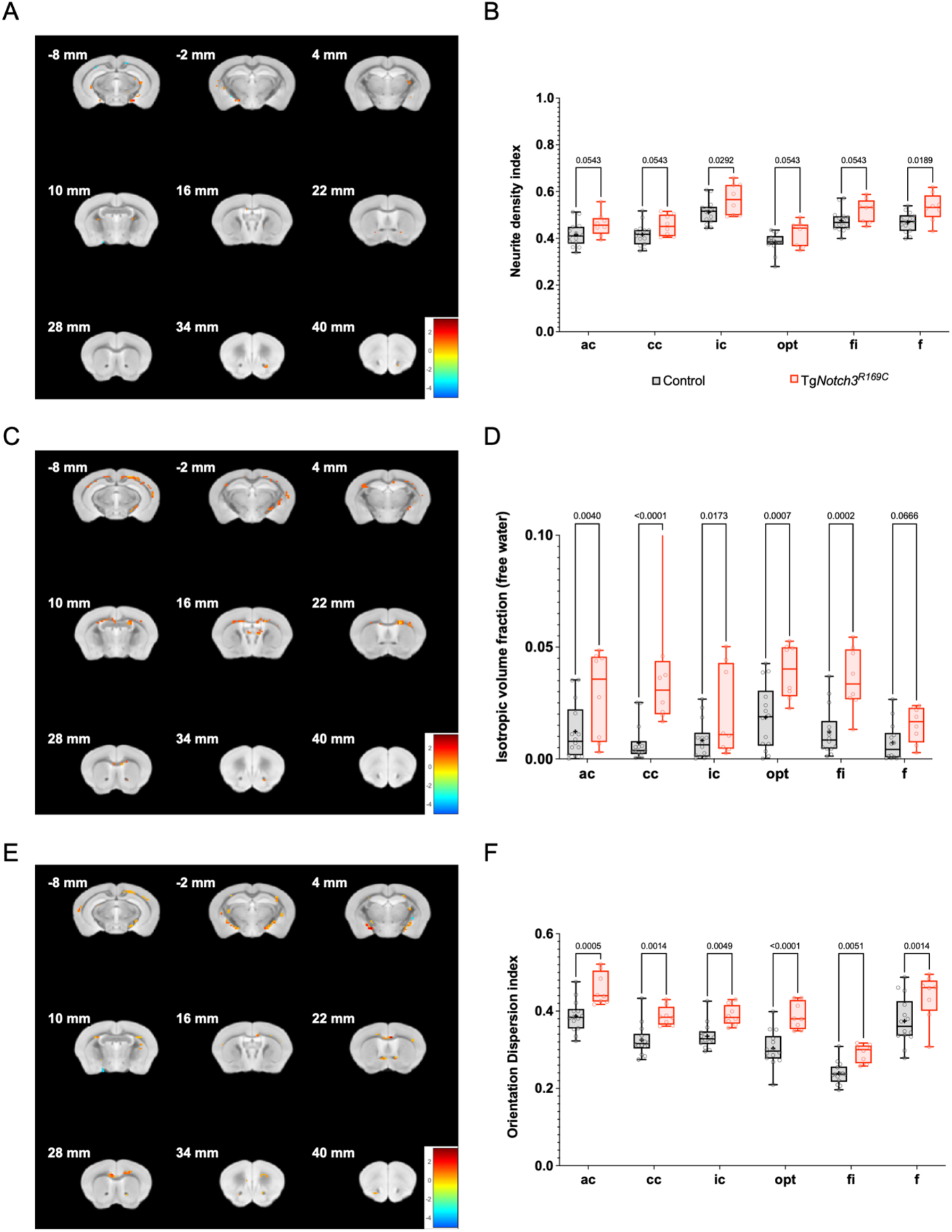
NODDI metrics in WM tracts of control and CADASIL mice. (**A, C, E**) Differences in voxel-based NDI (**A**), ISOVF (**C**) and ODI (**E**) values in WM tracts between Tg*Notch3*^R169C^ and control mice. (**B, D, F**) Box-and-whiskers plots of quantitative values in the specified WM tracts of Tg*Notch3*^R169C^ and control mice analyzed by a two-way ANOVA followed by a two-stage linear step-up procedure of Benjamini, Krieger and Yekutieli to correct for multiple comparisons. Shown are the q values. ac, anterior commissura; cc, corpus callosum; ic, internal capsula; opt, optic tract; fi, fimbria of the hippocampus and f, fornix.

**Figure 5:**
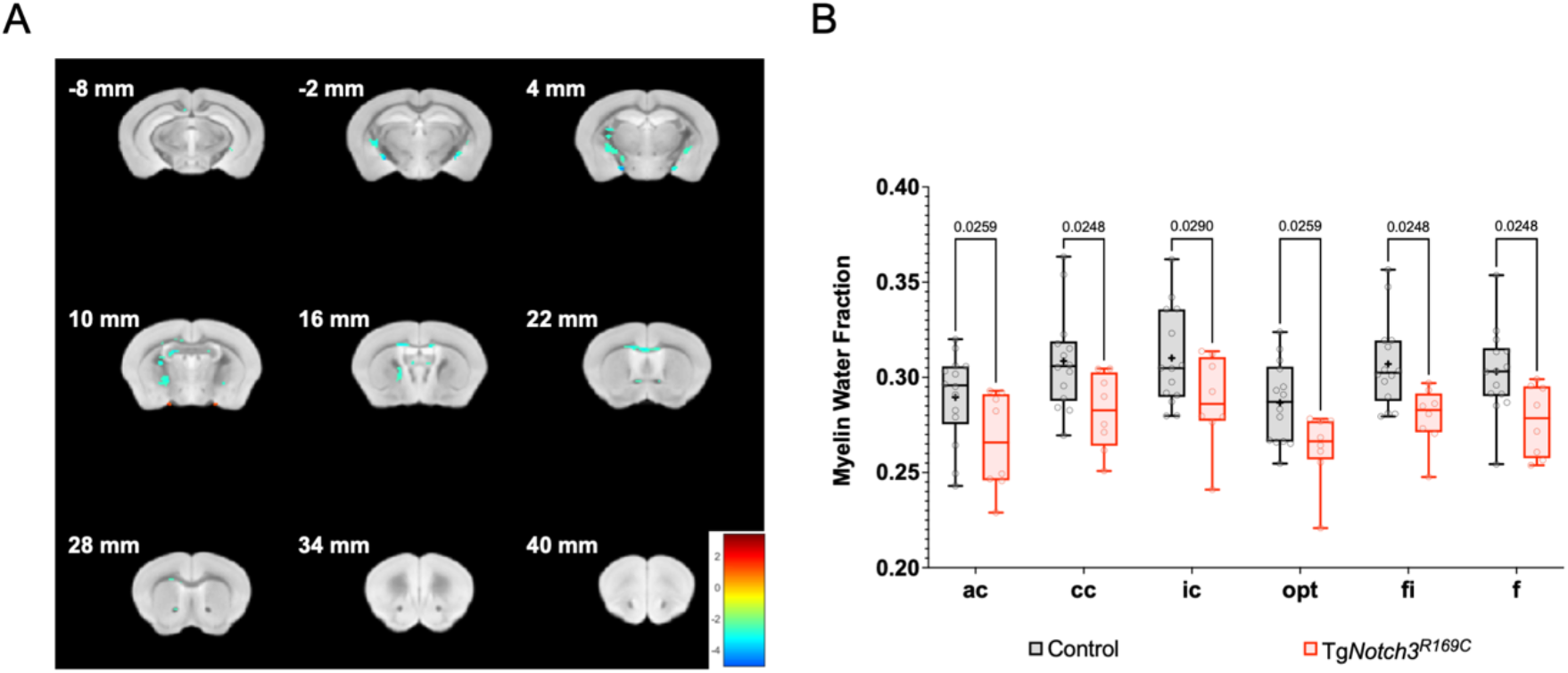
The MWF is widely reduced in hemispheric WM tracts of CADASIL mice. **(A)** Differences in voxel-based MWF values between Tg*Notch3*^R169C^ and control mice. (**B**) Box-and-whiskers plots of quantitative data in the specified WM tracts of Tg*Notch3*^R169C^ and control mice analyzed by a two-way ANOVA followed by a two-stage linear step-up procedure of Benjamini, Krieger and Yekutieli to correct for multiple comparisons. Shown are the q values. ac, anterior commissura; cc, corpus callosum; ic, internal capsula; opt, optic tract; fi, fimbria of the hippocampus and f, fornix.

**Figure 6:**
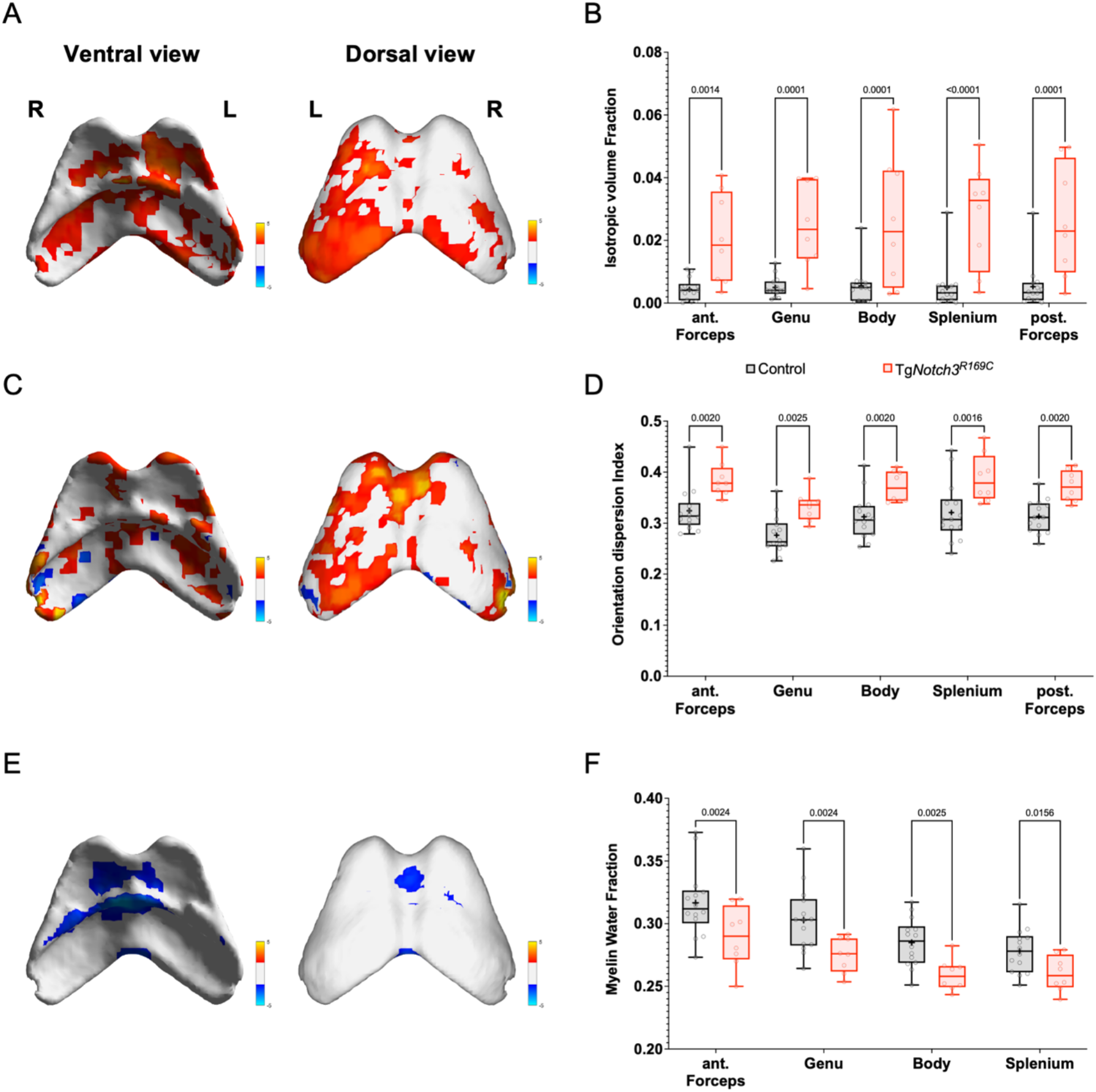
NODDI and MWF metrics in the different regions of the corpus callosum of control and CADASIL mice. (**A, C, E**) Differences in voxel-based ISOVF (**A**), ODI (**C**) and MWF (**E**) values in the corpus callosum between Tg*Notch3*^R169C^ and control mice. (**B, D, F**) Box-and-whiskers plots of quantitative data in the specified regions of the corpus callosum.

### Ultrastructural analysis

To assess the validity of our MRI findings, we performed electron microscopy analysis on the genu of the corpus callosum of Tg*Notch3*^R169C^ and control mice aged 22 months (Fig 7A). We used a second batch of mice because tissue preparation needed for high pressure freezing was radically different from the one used for MRI. We found that the percentage of myelinated axons was comparable between Tg*Notch3*^R169C^ and control mice (Fig 7B). However, the scatter plot of the g-ratio, a measure of the thickness of the myelin sheath relative to the axon diameter, showed an upshift of the cloud in Tg*Notch3*^R169C^ mice, indicative of myelin sheath thinning (Fig 7C). The large increase in extracellular free water detected in mutant mice (ISOVF; Fig 4C-D) prompted us to measure the extracellular spaces on electron micrograph images. In control mice, the extracellular space consisted of narrow slits between fibers (Fig 7A, left). In contrast, these spaces were wider in mutant mice (Fig 7A, right) and quantitative analysis revealed an almost two-fold increase in mutant mice compared with control mice (Fig 7D).

**Figure 7:**
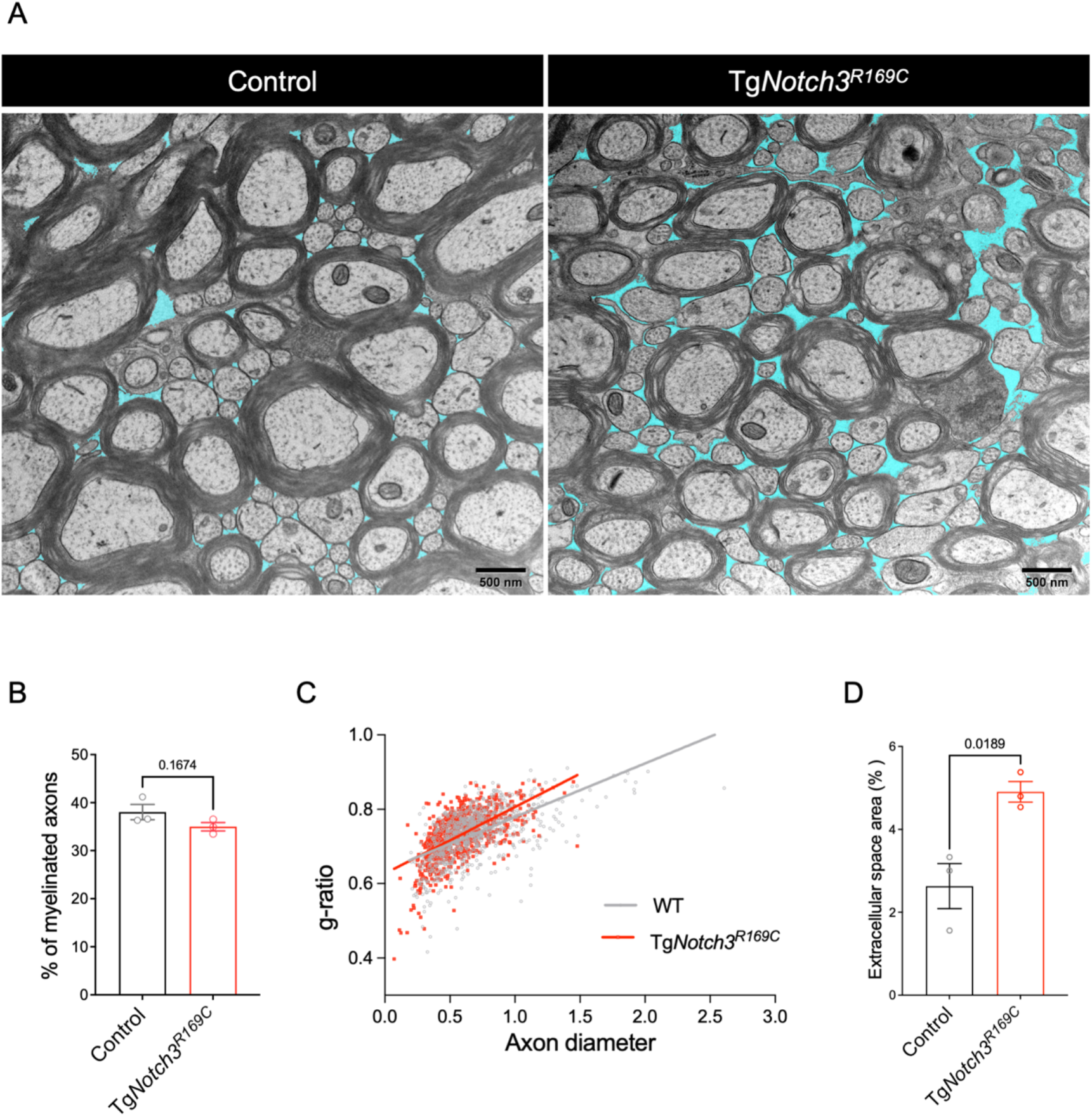
Myelin thinning and enlarged extracellular spaces in the corpus callosum of CADASIL mice. (**A**) Representative electron micrographs of corpus callosum samples (genu) from control (left) and Tg*Notch3*^R169C^ mice aged 22 months (right). Extracellular spaces are pseudo-colored in cyan. (**B**) Graph showing that the percentage of myelinated axons is comparable between control and CADASIL mice. (**C**) Plot of the g-ratio against axon diameter showing that the myelin sheath is thinner (higher g-ratio) in Tg*Notch3*^R169C^ mice. (**D**) Quantification of extracellular spaces showing an enlargement in Tg*Notch3*^R169C^ mice compared with control mice. Scale bars = 500 nm (A).

## Discussion

This is the first study combining dMRI and qMRI relaxometry with advanced multicompartment microstructural models to explore the pathological underpinnings of early WM signal abnormalities in the context of SVDs. Importantly, by applying these approaches to a SVD mouse model, this enabled us to confirm our MRI findings by ultrastructural analysis. We showed that GFA was widely reduced in the major hemispheric WM tracts of mice with CADASIL. We found that WM in mutant mice displayed a large increase in extracellular free water and a reduction in the myelin content, yet without reduction in neurite density. Remarkably, there was a high consistency between our MRI and histopathological findings. Indeed, electron microscopy analysis showed a ∽2-fold increase in the size of extracellular spaces and an elevation of the g-ratio indicative of myelin sheath thinning.

The use of ultra-high-field (11.7T) diffusion and qMRI in the frame of a prolonged acquisition protocol lasting 3 consecutive days allowed us to collect for each individual an extended set of dMRI volumes at the mesoscale level, at very high diffusion sensitizations (8000s/mm^2^) and with many diffusion directions (175 in total) along with an extended set of qMRI relaxometric volumes with an unusual large number of inversion and echo times. The scanning of each animal in a single session has eliminated the need to remove the sample from the scanner, thus avoiding the risk of motion artifacts between the acquisition of the different datasets. Combining dMRI and qMRI is a key point that has allowed us to accurately characterize the cytoarchitecture and myeloarchitecture of WM tracts at the voxel level. Another strength of our study was our ability to validate our MRI findings with high resolution electron microscopy, which is the gold standard technique to assess axonal and myelin integrity. Specifically, we used high pressure freezing and freeze substitution to process the samples, in order to better preserve the ultrastructure of WM fibers, which is a notorious issue in very old animals ^22^.

We recently reported that CADASIL mice exhibit myelin damages characterized by an enlargement of the inner tongue, and a splitting and thinning of myelin layers ^22^. A major new finding of this study is the marked increase in extracellular free water and extracellular space in old CADASIL mutant mice. The extracellular space is filled by interstitial fluid and its volume fraction with respect to the total brain is reported to be around 20%, although there could be regional variations ^23^. In our ultrastructural studies the volume or space fraction in control animals was much lower because we used aldehyde fixation prior to high pressure and freeze substitution of samples. Fixatives are known to shrink extracellular spaces, likely because of an imbalance in the osmolarity across cellular membranes ^24^. However, since control and mutant brain samples have been processed similarly, we are confident that this large increase in extracellular free water and extracellular space in mutant mice is real and not artefactual. Interestingly, two recent studies applied free water imaging to SVD patients including young CADASIL patients and found increased extracellular free water as the main source of diffusion alterations in patients ^25,26^. In these two studies, free water measurement was derived from dMRI using a 2-compartment model based on second order tensors, including one compartment representing water molecules that are not restricted and assumed to be extracellular water and a second tissue compartment representing all remaining water molecules. Herein, the free water compartment was derived from the multicompartment NODDI model, through the ISOVF parameter, in a more realistic and robust way than the free water corrected tensor model mentioned above since it accounts for a better mathematical solution of the diffusion process occurring within the axonal space using sticks to represent axons.

Our results argue against the possibility that interstitial fluid accumulation and enlarged extracellular spaces in CADASIL mutant mice are merely the consequence of a reduction in axonal density. Moreover, extracellular spaces were measured on electron micrographs of WM tracts away from vessels, and thus were clearly distinct from perivascular spaces whose enlargement has been repeatedly reported in SVD patients, and especially in CADASIL patients ^27^. Taken together these results suggest that accumulation of interstitial fluid is a major and early pathological process in SVD. The relationship between this accumulation of fluid and myelin sheath damages remains to be explored.

Several mechanisms could lead to interstitial fluid accumulation in the WM. One possible explanation is an increased permeability of the blood brain barrier (BBB). In the present CADASIL mouse model, we did not find evidence of leakage of endogenous and exogenous tracers into the brain parenchyma ^28^. Also, Markus and colleagues did not detect an increased BBB permeability to Gadolinium in CADASIL patients ^29^. However, Yang and colleagues recently reported a reduction in the water exchange rate across the BBB in patients with CADASIL or HTRA1-related SVD, as measured by pseudo-continuous arterial spin labeling, suggestive of an increased permeability of the BBB to water ^30^. Hence, CADASIL could be associated with a dysfunctional BBB predominating on water and ion permeability, at least at the early stage of the disease. Another attractive possibility is a defective drainage of interstitial fluid due to pathological changes in the glymphatic system. This system transports extracellular fluid into the brain along and through the arterial perivascular spaces and out of the brain via the perivenous spaces and is driven by many of the same processes that regulate brain perfusion ^31^. Any imbalance between the glymphatic influx and the glymphatic efflux is expected to cause interstitial fluid accumulation and extracellular spaces enlargement. In support of a possible failure of brain fluid transport in SVDs is the striking enlargement of perivascular spaces, which are the main route for the drainage of brain fluid ^27,32^.

We acknowledge some limitations of our work including the cross-sectional design of our study and the MRI imaging of post-mortem brains rather than alive animals. However, this study was designed as a proof-of-concept aiming at achieving the highest possible spatial resolution. Aldehyde fixation of brain samples in our study has caused us to underestimate the volume of extracellular space and extracellular free water, however using comparable aldehyde fixation for MRI and electron microscopy analyses enabled cross-validation between the two approaches.

In summary, our findings provide compelling evidence that accumulation of interstitial fluid and myelin sheath damage are 2 major factors underlying early WM changes in CADASIL. Moreover, our work indicates that a multimodality imaging approach, combining multi-shell dMRI, relaxometry and advanced biophysical modeling is highly powerful to decipher the pathological underpinnings of WM lesions in SVD. Determining axonal and myelin integrity as well as extracellular free water content should provide better insight into differentiating the severity of lesions that appear similar on conventional MRI imaging, and allow better monitoring of disease progression in patients. Though our findings are in a preclinical model, translating these into a clinical setting should be relatively straightforward as most MRI techniques for small animals are the same as those already used in clinics.

## Acknowledgements

We are grateful to Xavier Heiligenstein and Ilse Hurbain for technical assistance with the HPF/FS from the cell and tissue imaging facility (PICT) of Institut Curie.

## Author contributions

DB and IU performed the MRI analysis and analyzed the data, DB wrote a first draft of the manuscript. RMR, assisted by VDD and FG, conducted the ultrastructural study and analyzed the data. SM, FB and JB assisted with the MRI experiments. CP & AJ designed the study, supervised research, analyzed data and wrote the manuscript.

## Data Availability Statement

The data that support the findings of this study and the codes are available from the corresponding author upon reasonable request.

## List of abbreviations

BBB: blood brain barrier
CSF: cerebrospinal fluid
dMRI: diffusion MRI
DTI: Diffusion tensor imaging
FOV: field of view
GFA: Generalized Fractional Anisotropy
ISOVF: Isotropic volume fraction
MRI: magnetic resonance imaging
MWF: Myelin water fraction
NODDI: Neurite orientation dispersion and density imaging
NDI: neurite density index
ODI: orientation dispersion index
qMRI: quantitative MRI
SVD: small vessel disease
TE: echo time
TR: repetition time
WM: white matter

